# Integrating habitat suitability modeling with gene flow improves delineation of landscape connections among African savanna elephants

**DOI:** 10.1101/2023.08.22.554325

**Authors:** Alida de Flamingh, Nathan Alexander, Tolulope I.N. Perrin-Stowe, Cassidy Donnelly, Robert A.R. Guldemondt, Robert L. Schooley, Rudi J. van Aarde, Alfred L. Roca

## Abstract

Across Africa, space for conservation is sometimes limited to formally protected areas that have become progressively more isolated. There is a need for targeted conservation initiatives such as the demarcation of landscape connections, defined as areas that encompass environmental variables that promote the natural movement of individuals between populations, which can facilitate gene flow. Landscape connections can mitigate genetic isolation, genetic drift, and inbreeding, which can occur in isolated populations in protected areas. Promoting gene flow can reduce the risk of extirpation often associated with isolated populations. Here we develop and test models for identifying landscape connections among African savannah elephant (*Loxodonta africana*) populations by combining habitat suitability modeling with gene flow estimates across a large region including seven countries. We find a pronounced non-linear response to unsuitable habitat, consistent with previous studies showing that non-transformed habitat models are poor predictors of gene flow. We generated a landscape connections map that considers both suitable habitats based on telemetry occurrence data and gene flow estimated as the inverse of individual genetic distance, delineating areas that are important for maintaining elephant population connectivity. Our approach represents a novel framework for developing spatially and genetically informed conservation strategies for elephants and many other taxa distributed across heterogeneous and fragmented landscapes.

**GRAPHICAL ABSTRACT:** 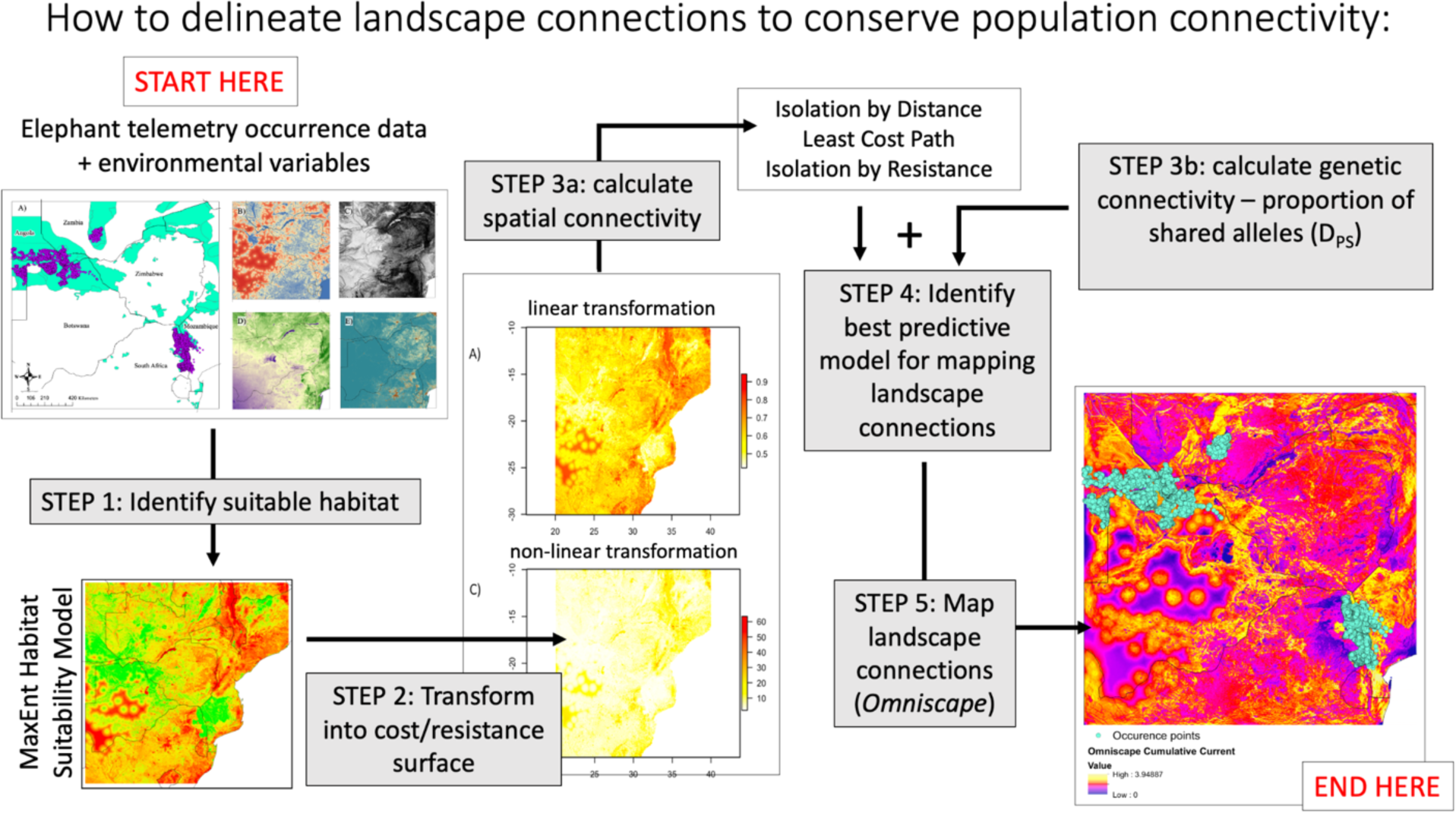

## INTRODUCTION

Across much of Africa, space for conservation may be limited to formally protected areas that have become progressively more isolated (Newmark, 2008; Ripple et al., 2015). However, despite the compression of African elephants (genus *Loxodonta*) populations into these fragmented protected areas and the resulting potential isolation of elephant populations into discontinuous units, southern African elephant (*Loxodonta africana)* populations do not yet demonstrate characteristics of genetically isolated populations (Garant et al., 2007; Allendorf et al., 2013). Instead, many southern African populations demonstrate genetic diversity that is similar to or higher than populations in east Africa (Lobora et al., 2018; de Flamingh et al., 2023). The genetic diversity of southern African populations has likely been maintained due to the large size of the surviving elephant populations, long elephant generation times, and dispersal and gene flow across the region (de Flamingh et al., 2018). However, there may have been historical connections that are no longer good candidates for connectivity restoration especially with dynamic landscapes or anthropogenic land conversion (Huang et al., 2022). Given the large home range of elephants, connectivity analyses can help identify effective conservation strategies within both protected and unprotected areas to support population persistence by increasing dispersal, gene flow, genetic diversity, and environmental adaptability (Garant et al., 2007).

To understand how gene flow can be maintained despite habitat fragmentation, we focus on landscape connections, or areas that encompass environmental variables that promote the natural movement of individuals between populations. These connections would facilitate gene flow and thereby minimize the genetic isolation and inbreeding that can occur in isolated populations in protected areas. Thus, landscape connections can be instrumental in increasing the genetic health and the persistence of wildlife populations (Allendorf et al., 2013). Landscape connections may be more robust when identified by integrating spatial and genetic data, rather than relying only a single data type. Spatial data may demonstrate habitat suitability and potential routes for dispersal but would not provide evidence of gene flow across populations. By contrast, genetic data may establish that gene flow has occurred among populations but would not identify the specific geographic routes needed to maintain it.

To integrate these data, landscape genetics merges aspects of population genetics with landscape ecology, providing a useful framework to identify gene flow (Manel et al., 2003; Manel and Holderegger, 2013). However, methodology and applications vary widely (Riordan-Short et al., 2023). Often expert opinion is used to identify barriers to gene flow, but this method is criticized for non-empiricism (Spear et al., 2010; Milanesi et al., 2017). Habitat suitability models offer a more objective approach (Milanesi et al., 2017), but may be inappropriate for modeling connectivity during specific life events like mating and natal dispersal (Keeley et al., 2017). Furthermore, the spatial scale of habitat suitability models may vary, and landscape genetic studies may use habitat use within a home range (e.g., Shafer et al., 2012; Keeley et al., 2017)or range-wide distribution HSMs to parameterize models (e.g., Shrestha and Kindlmann, 2020), which are different orders of habitat selection (Johnson, 1980). Species may interact differently with the landscape when within home ranges than during dispersal events (Centeno-Cuadros et al., 2011; Büchi and Vuilleumier, 2014; Alexander et al., 2019).

In this study, we create a habitat suitability map, visualize spatial genetic patterns across the study area, and then estimate gene flow and connectivity using a landscape genetic approach. We then delineate landscape connections among African elephant populations in seven southern African countries. We parameterized a commonly used presence-only habitat suitability model, MaxEnt (Philips et al. 2006, 2017). Presence-only models can be expanded to integrate presence-absence data (Renner and Warton, 2013; Fletcher et al., 2016; Koshkina et al., 2017), positioning this established HSM framework to be a robust method to empirically identify landscape connections. Next, we created a genetic landscape through a graph-theory network approach (Miller 2005). Finally, we assessed how the top HSM, and logarithmic transformations of the top HSM, correlate to gene flow and identify landscape connections. Overall, we apply this approach to identify suitable habitat patches, spatial genetic structure, and how gene flow is related to habitat suitability for the African savannah elephant, an endangered, wide-ranging mammal (Gobush et al., 2022).

## METHODS

Although understanding species distributions are important for conservation, it is also critical to understand how landscape features impact gene flow and produce spatial genetic patterns (Manel et al., 2003; Holderegger and Wagner, 2008; Richardson et al., 2016). Habitat suitability models can be useful as an empirical approach to understanding which landscape features impact species distribution (Spear et al., 2010; Milanesi et al., 2017). We used the predicted potential species distribution from MaxEnt to create a habitat suitability map (HSM) that we transformed linearly and non-linearly into resistance surfaces (Keeley et al., 2016). These transformations invert the HSM values so that areas of high suitability have low resistance values (Keeley et al., 2016). We considered nonlinear transformations to investigate possible threshold responses that elephants might have to unsuitable habitats in the resistance surfaces. Landscape resistance surfaces are grid maps that represent the resistance to traversing the landscape features contained in that grid cell (Milanesi et al., 2017), or the cost of moving through such a landscape. To evaluate alternative quantifications of connectivity across the landscape, we fitted maximum likelihood population effects (MLPE) models (Clarke et al., 2002; Van Strien et al., 2012) between pairwise matrices of genetic distances and matrices of Euclidean, least cost path (LCP) or circuit theory (CT) distances (Shah and McRae, 2008). Our MLPE model assessment identified CT distances as the connectivity measure that best explained the estimated gene flow across our landscape. We then used Omniscape to generate connectivity maps across the study region (McRae et al., 2016; Landau et al., 2021). Omniscape estimates connectivity using circuit theory (McRae, 2006; McRae and Beier, 2007; Shah and McRae, 2008) and a moving window approach to generate current maps, or an estimate of the number of random-walks that pass through a location, that are then summed to create a landscape-scale connectivity map. Our final map therefore considered suitable habitats based on spatial data, and also gene flow as the inverse of genetic distance, to delineate landscape connections that should be important for maintaining population connectivity for elephants.

### Study area and elephant occurrence data

Our study area spans seven countries across southern Africa, including Angola, Botswana, Mozambique, Namibia, South Africa, Zambia and Zimbabwe (Figure 1), and includes a range of different vegetation classes (SI Figure 1). Together these countries contain >70% of the elephants of Africa and >42% of the total range of elephants in Africa (Thouless et al., 2016). Elephants occur across landscapes that have a range of precipitation and vegetation characteristics (Loarie et al., 2009a; Roever et al., 2013). These include, but are not limited to, areas of very low precipitation such as desert landscapes in Namibia (Ishida et al., 2016), to areas with intermediate precipitation such as wetland landscapes in the Okavango Delta of Botswana (Songhurst et al., 2015), to high precipitation areas such as the mesic savannas of Malawi and Zambia (Ott, 2008) and the forest and Futi floodplains of Maputo Special Reserve in Mozambique (Ntumi et al., 2005).

**Figure 1.**
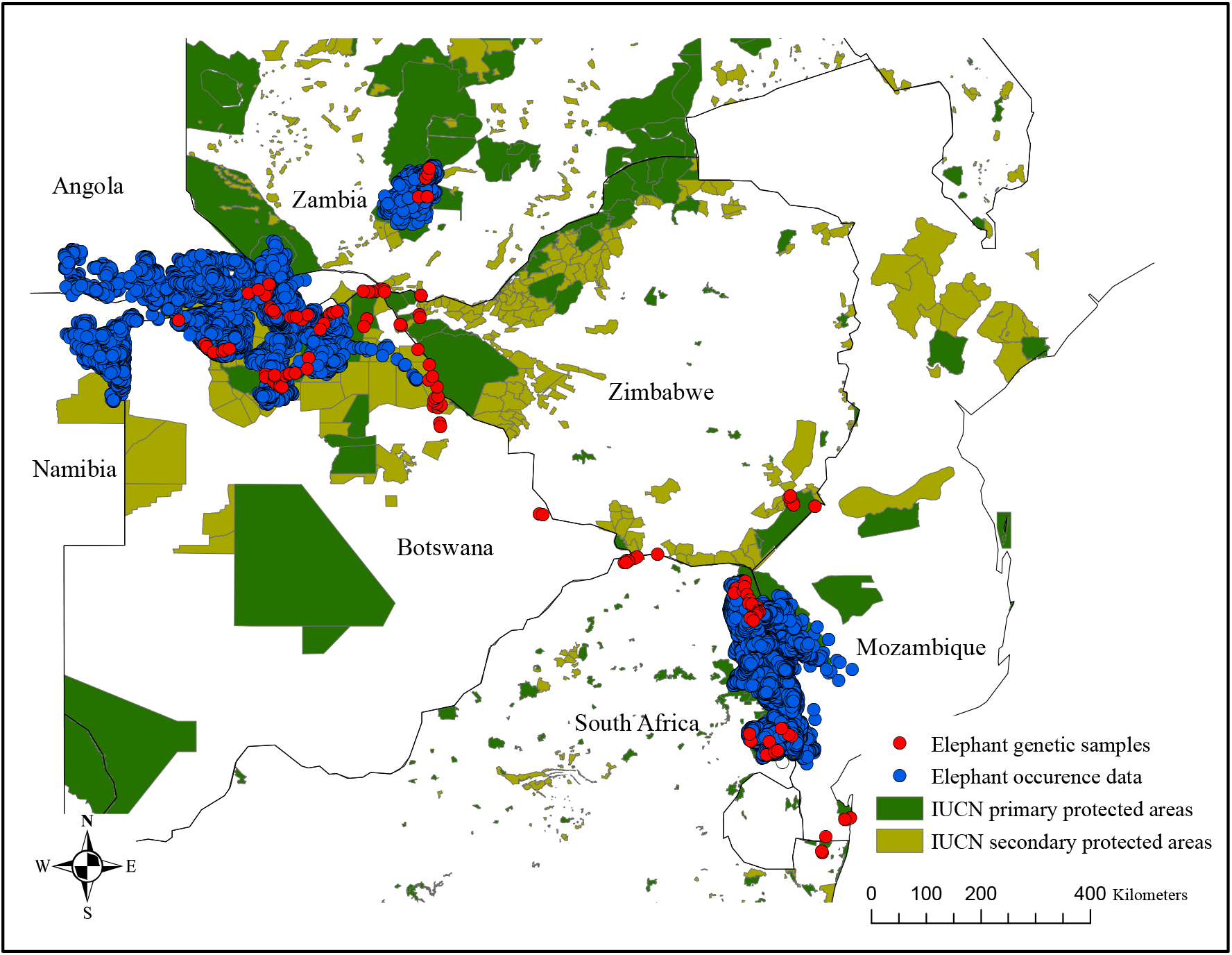
Map of the study area spanning seven countries across southern Africa, including Angola, Botswana, Mozambique, Namibia, South Africa, Zambia and Zimbabwe. These countries together contain >70% of the number of elephants in Africa and >42% of the total range of elephants in Africa (Thouless et al., 2016). Occurrence data (GPS telemetry points) from 80 elephants are shown in blue, and locations at which genetic samples were collected are shown in red.

The Conservation Ecology Research Unit (University of Pretoria, South Africa) has an extensive database of elephant telemetry (GPS collar) data (Figure 1 – Elephant occurrence data; Figure 2A). We used quality filtering criteria similar to Roever et al. (2013) to select reliable telemetry locations. To decrease temporal and spatial autocorrelation, we filtered the data so that only a single point per day per elephant was retained. Our final occurrence dataset comprised 80 elephants (15 male and 65 female) and 53,954 location points. These elephants fall within three regional population clusters that include the Chobe, Kafue, and Limpopo clusters (SI Figure 2).

**Figure 2.**
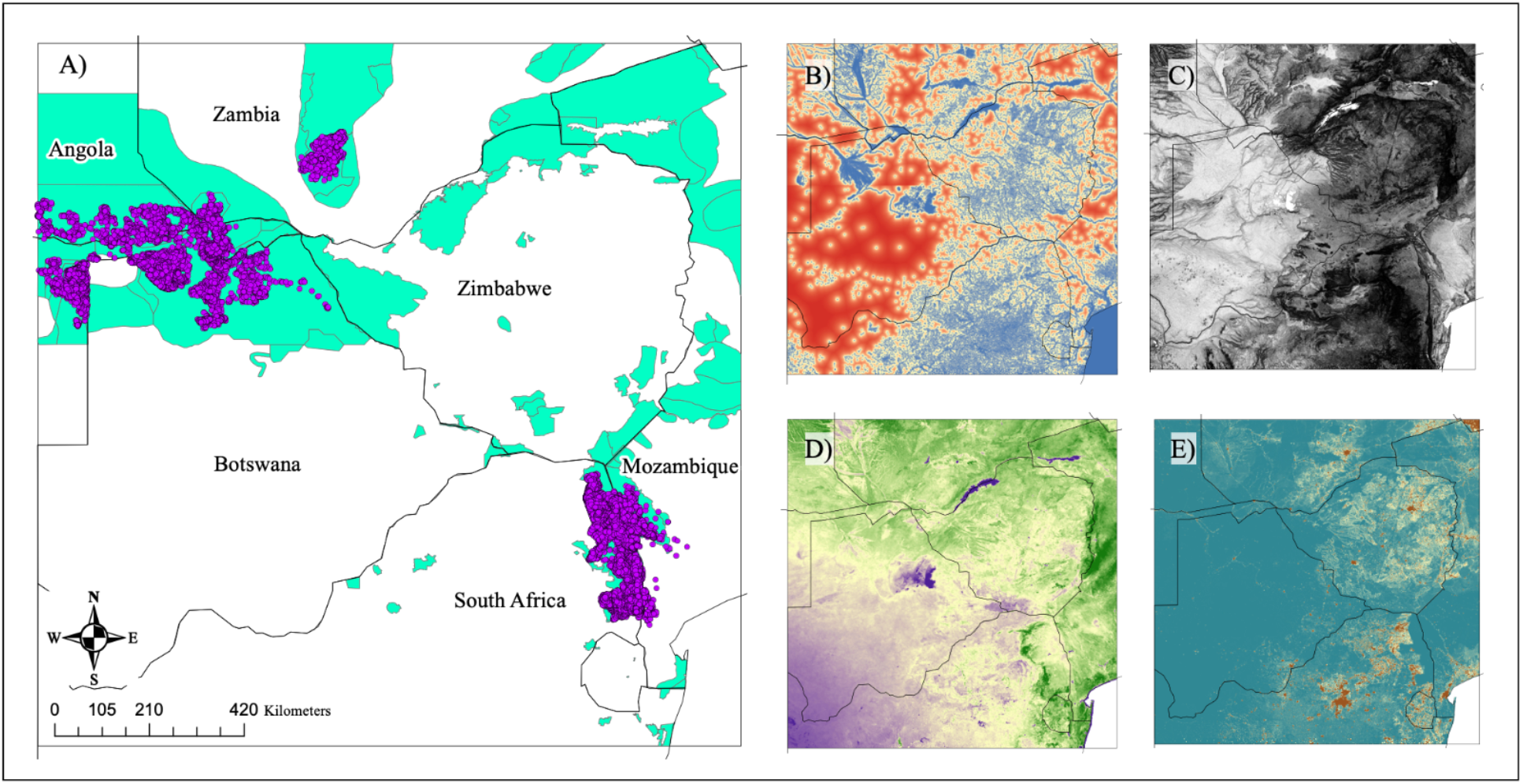
The top MaxEnt habitat suitability model was based on elephant occurrence points that correspond with areas of known elephant range (panel A - purple dots represent occurrence data and green areas represent the current elephant range as demarcated by the IUCN Red List of Threatened Species; Blanc, 2008). MaxEnt environmental variables included habitat covariates that are known to influence elephant space use: water availability as represented by distance to the closest water source (panel B – blue areas are close to water, red areas are far from water), the slope of the land (panel C – dark areas represent steep slopes), primary productivity indexed by long-term mean Enhanced Vegetation Index (EVI; panel D – green areas indicate high primary productivity, brown and purple areas indicate low primary productivity), and ambient human population density (panel E – red indicates areas with high ambient population density, blue indicates areas with low ambient population density).

### Elephant genetic data

Fresh elephant fecal samples were collected from six southern African countries from 2010-2014 (Figure 1 – Elephant genetic samples). A total of 142 samples were collected and genotyped for 9 highly variable nuclear DNA microsatellite loci (de Flamingh et al., (2018); SI Table 1). Sample collection, DNA extraction procedures, and microsatellite amplification are detailed in de Flamingh et al. (2018), and genotype errors were quantified as described in de Flamingh et al. (2014). Heterozygous genotypes were replicated at least four times, and homozygous genotypes were replicated at least three times. Diversity indices for individual microsatellite loci were calculated in Arlequin (Excoffier and Lischer, 2010)

**Table 1.** MaxEnt models that included background data (n = 10,000) from the entire geographic extent of our study area had the highest area under the receiver-operator curve (AUC), while the range-based background dataset resulted in slightly higher AUC than use-based background dataset.

| Background data | Maxent regularization parameter | Regularized training gain | AUC |
| --- | --- | --- | --- |
| *Range-based background data\** | $\beta = 1$ | 0.039 | **0.786** |
| | $\beta = 2$ | 0.034 | 0.781 |
| | $\beta = 3$ | 0.030 | 0.775 |
| | $\beta = 4$ | 0.027 | 0.764 |
| *Use-based background data* | $\beta = 1$ | 0.039 | 0.758 |
| | $\beta = 2$ | 0.035 | 0.755 |
| | $\beta = 3$ | 0.032 | 0.751 |
| | $\beta = 4$ | 0.030 | 0.745 |

\*The top MaxEnt model that was calculated using the range-based background dataset and a regularization parameter of $\beta = 1$ for subsequent analyses because it outperformed the use-based dataset.

### Habitat suitability modeling

#### Parameter selection

We considered habitat covariates that are known to influence the distribution of elephants including water availability (Loarie et al., 2009b), the slope of the land (Wall et al., 2006), primary productivity (Young et al., 2009a) and human population distribution (Hoare and Du Toit, 1999). Surface-water availability drives the distribution and abundance of elephants (Chamaillé-Jammes et al., 2007). Elephants prefer areas that are close to water, especially during the dry season when access to surface water is limited. Elephants alter their space use to avoid mountainous terrain, as even small hills may present energetic barriers to movement for large-bodied animals (Wall et al., 2006). Primary productivity as a proxy for food availability influences elephant habitat selection and space use (Young et al., 2009b; Roever et al., 2012). In addition, elephants also alter their space use to avoid areas densely populated by humans (Barnes et al., 1991; Hoare and Du Toit, 1999). These four environmental variables included in this study thus directly impact elephant space use and the potential geographic distribution of landscape connections.

We converted the Global Surface Water (GSW) occurrence layer (Pekel et al., 2016) to estimate distance to water across our study range. The GSW occurrence layer represents areas where surface water occurred between 1984 and 2018 and provides information concerning overall water dynamics. This layer captures the frequency with which water was present in a given area. To calculate GSW occurrence, Pekel et al. (2016) respectively summed and then normalized by division the monthly water detections (WD) and valid observations (VO) such that GSW occurrence/month = ∑WD month / ∑VO month. By averaging the results of all monthly GSW occurrence calculations, Pekel et al. (2016) were able to provide the long-term overall surface water occurrence.

GSW data were downloaded as 9 individual tiles that covered our study extent and were merged using the “Mosaic to new raster” function in ArcMap V10.7.1 (© ESRI 2011). We transformed the GSW data to represent the geographic distance from an available water source because areas that are further from water sources would represent less suitable habitat, and a gradient layer may therefore be more biologically relevant to elephant space use than a binary presence-absence layer. We reclassified the GSW data so that all points with values >0 were reclassed to represent water occurrence. We used the Euclidean distance tool in ArcMap to calculate, for each cell in our study area, the Euclidean distance to the closest water source (Figure 2B). The GSW dataset was split into subset of four raster layers before executing the Euclidean distance tool to facilitate data processing.

We converted a digital elevation model (Jarvis et al., 2008) to denote slope by calculating the maximum rate of elevation change between pixels using the “Spatial Analyst” toolbox in ArcMap (ESRI © 2011). We use this converted slope layer as our second environmental layer in MaxEnt (Figure 2C).

We used long-term mean Enhanced Vegetation Index (EVI) from Robson et al. (2017), which is an index of primary productivity across our study range (Figure 2D). We use EVI rather than Normalized Difference Vegetation Index (NDVI) because EVI overcomes some of the contamination problems present in NDVI data (e.g., contamination associated with canopy background and residual aerosol influences), and EVI is less likely to become saturated in areas that have high green biomass (Pettorelli et al., 2005).

We used Landscan2016 global population distribution data, which is the finest resolution human population data available. Landscan data represents an “ambient population” (average population distribution over 24 hours; Bright et al., 2017). LandScan data are preferable to other census-based data because they account for differences in spatial data availability, scale and accuracy, and Landscan distribution models are specific to individual countries and regions (Bright et al., 2017). Furthermore, LandScan data as an “ambient population” rather than point density consider both diurnal movements and collective travel habits and integrates these variables into a single measure (Dobson et al., 2000). Other studies have transformed human density data to a gradient-based distance metric that represents geographic distances from densely populated areas (Roever et al., 2013). We assessed both transformed, gradient-based human density and raw LandScan “ambient population” data in our habitat suitability models (Figure 2E). Using raw ambient human population data may better align with a nonlinear relationship in which elephant and human coexistence occurs at a range of human densities, but local absences of elephants are contingent on landscape-dependent thresholds of human density (Hoare and Du Toit, 1999).

#### Raster preparation for habitat suitability modeling

MaxEnt requires all environmental layers to have the same resolution and spatial extent. We resampled environmental layers to 900m X 900m (0.0083 x 0.0083 decimal degrees; standardized across environmental data layers) so that resolution represented a biologically relevant scale in view of elephant space use and distribution. The spatial resolution of our layers is informative for a highly mobile species such as elephants (Young et al., 2009b) but is still much smaller than average home range size of elephants in these population clusters (Grainger et al., 2005; Roever et al., 2012). We used a bilinear interpolation approach that is suitable for continuous data to resample our environmental layers using the “Data management” toolbox in ArcMap (ESRI © 2011). Bilinear interpolation calculates the value of each pixel by averaging the values of surrounding pixels. We standardized the coordinate system and datum to WGS_1984 for all input layers and tested for correlation among our environmental variables by calculating Pearson’s correlation coefficient in the R package “virtualspecies” (Leroy et al., 2016; R Core Team, 2019)

#### Habitat suitability analysis

We predicted the potential distribution of elephants using a maximum-entropy (MaxEnt) species distribution modeling approach (Phillips et al., 2006, 2017). MaxEnt has been used extensively to predict the potential distribution of animals and is a widely used software for distribution modeling in conservation studies (Fourcade et al., 2014). MaxEnt moves beyond classical regression methods such as resource selection functions (Boyce and McDonald, 1999; Chetkiewicz and Boyce, 2009) and generalized linear models (Guisan et al., 2002) and employs an algorithmic modeling approach that is based on machine learning (Fourcade et al., 2014). MaxEnt estimates the potential distribution of animals by finding the spatial distribution of maximum entropy (the distribution that is closest to being uniform) when comparing the expected value of each environmental variable under this estimated distribution to the empirical average that has been calculated from the occurrence or telemetry data (Merow et al., 2013). MaxEnt is especially efficient when there is a complex relationship between response and predictor variables (Elith et al., 2011), and MaxEnt has outperformed other modeling approaches to predict animal distribution (Hernandez et al., 2006; Wisz et al., 2008; Shabani et al., 2016).

We used MaxEnt to determine which environments represented suitable habitats for elephants. MaxEnt requires presence-only data to model species distribution and predict habitat suitability (Elith et al., 2011). MaxEnt uses occurrence data points (presences; SI Figure 3A) in combination with a set of environmental predictor variables (e.g., distance to water, slope, vegetation index, human presence), and compares these presences and their associated environmental constraints to a set of random background points (Merow et al., 2013). MaxEnt then predicts the relative occurrence rate (ROR) for each cell in the landscape grid, where ROR is the relative probability that the cell is contained in the presence data, and where ROR is contingent on selecting the “raw” output option in MaxEnt (Elith et al., 2011). MaxEnt can also be used to estimate the probability of presence by using a logistic transformation of the ROR that allows for large differences in output values to translate to large differences in suitability. Logistic probability of presences may improve model calibration and are contingent on selecting the “logistic” output option in MaxEnt (Phillips and Dudík, 2008). We only compare map predictions that were generated using the same assumptions to estimate the probability of presence (Merow et al., 2013).

**Figure 3.**
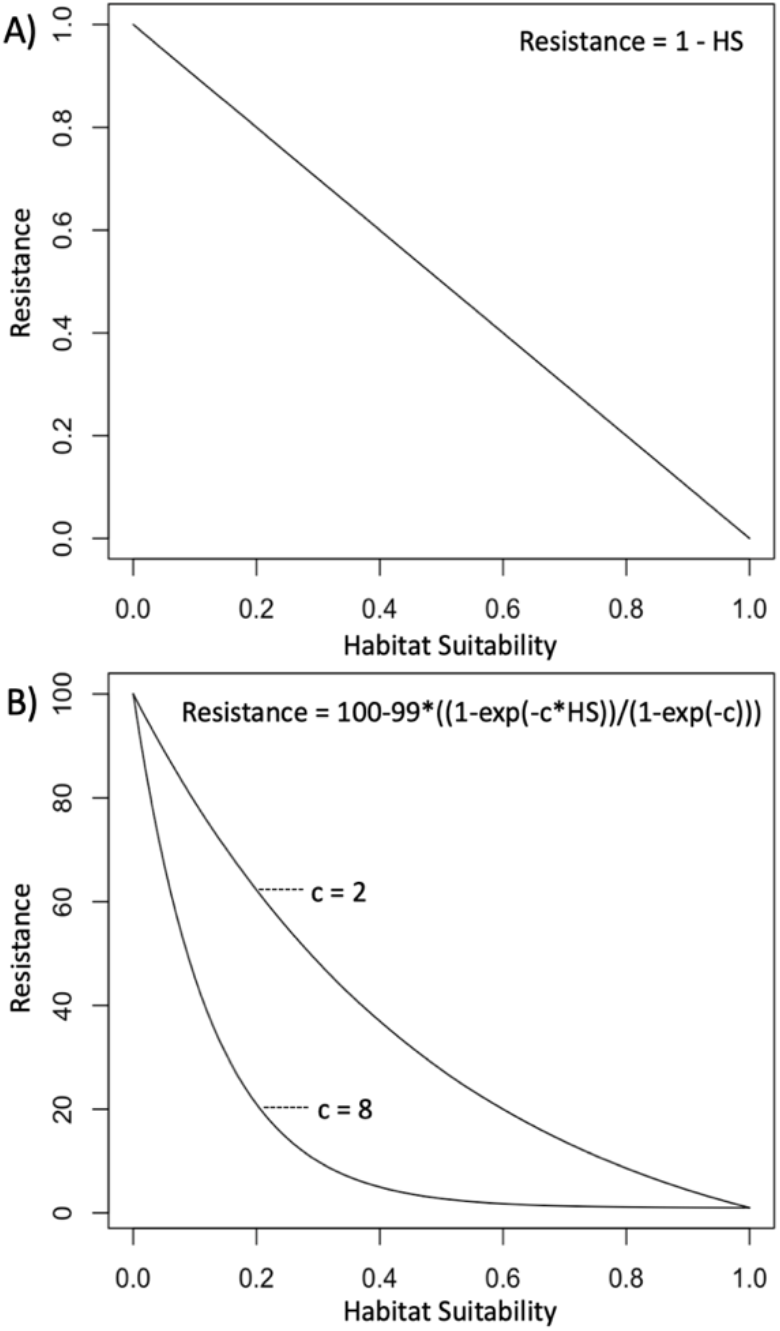
The single best habitat suitability model was transformed linearly (panel A) and nonlinearly (panel B) into resistance surfaces. A linear transformation assumes that resistance is inversely related to habitat suitability, where less suitable habitat would be costlier to move through. A transformation by means of a negative exponential function allowed for the possibility of nonlinear responses that elephants might have to unsuitable habitats. We generated resistance surfaces for transformations using c = 2 and c = 8 to respectively generate slight and pronounced nonlinear responses. Equations for linear and nonlinear transformations are provided in the upper right corner of each graph.

#### MaxEnt background data

Choosing relevant background location data for comparison to presence data is critical when the MaxEnt model will be used to predict or extrapolate to novel environments (Elith et al., 2011; Webber et al., 2011). Background data should be chosen to reflect the environmental conditions that are relevant to the species for which the model is generated, and they should be based on the spatial scale of the ecological questions of interest (Saupe et al., 2012; Merow et al., 2013; SI Figure 3). Model fit metrics can be inflated by increasing the number of background points, and by selecting background data that may not be ecologically informative for species distribution modeling (Lobo et al., 2008; Anderson, 2012).

We aimed to delineate landscape connections within and across areas where elephants are known to occur. We were also interested in delineating connections that may not currently fall within the known elephant range, but where future initiatives may aim to conserve these corridors to re-establish connectivity between isolated populations. We therefore tested two different background datasets: range-based background data generated for areas of known elephant range as demarcated by the IUCN Red List of Threatened Species (SI Figure 3B, Blanc, 2008); and use-based background data from areas included in an 80% convex hull of the presence points (SI Figure 3C). For each of these datasets, we generated 10,000 random points to compare to the presence data.

#### MaxEnt model regularization

Model regularization can reduce model complexity and over-fitting (Merow et al., 2013). Regularization may allow for less precise fitting of the empirical constraints from environmental features in the MaxEnt model, and it simplifies models by incorporating a penalty that is proportional to the magnitude of the regularization coefficient (Merow et al 2013). Therefore, using explicit regularization parameters may prevent model complexity from increasing beyond what is supported by the empirical dataset (Phillips and Dudík, 2008). We tested four regularization beta parameters in Maxent (*β* = 1, 2, 3, 4), and compared the area under the receiver-operator curve (AUC) values for cross-validated models as a measure of model fit to determine the most suitable beta parameter for our dataset. Regularization parameters were tested for both background datasets (Table 1).

#### Model evaluation

We evaluated the Maxent models using a 5 k-fold cross validation. We withheld a random sample of 20% of the presence data for testing and trained our model with the remaining 80% of the data. We assessed the evaluated models by investigating the AUC. AUC calculates the probability that a presence location ranks higher than a random background point in the predicted model (Merow et al., 2013). We compared AUC values for both types of background data, where each type of background data was fitted using *β* = 1, 2, 3, 4 (8 models in total). We selected the best scoring combination of background type and regularization parameter to generate our HSM.

### Spatial visualization of gene flow

To visualize gene flow across the landscape, we interpolated genetic distances to form a landscape shape in the program *Alleles In Space* (AIS; Miller, 2005). AIS creates a connectivity network across all sample locations and places genetic distances as midpoints of each pairwise connection. The program then interpolates genetic distances across the extent of the study area and produces a 3-dimensional surface plot in which surface heights represent interpolated genetic distances, and where higher peaks in the surface plot indicate greater genetic distances. We tested for correlation between genetic and geographic distances, and calculated genetic distance in AIS as in Nei et al. (1983). We used a distance weighting parameter (*a*) of 1 to interpolate genetic distance across the landscape (Miller, 2005). We exported the interpolated output using the highest possible resolution of 500 x 500 bins for the X and Y geographic axes, and plotted for each of the bin coordinates the peak heights using ArcMap V10.7.1 (© ESRI 2011).

### Landscape genetics

We transformed the top HSM, linearly and nonlinearly, through a negative exponential function into resistance surfaces (Keeley et al., 2016). Resistance is usually assumed to be a negative linear function of suitability (Hunter et al., 2003; Larkin et al., 2004; Pullinger and Johnson, 2010). However, in addition to a negative linear transformation, we also tested slight and pronounced nonlinear transformations to investigate potential threshold responses by elephants to unsuitable habitats in the resistance surfaces. A linear transformation assumes that cost (R_lin_) is inversely related to habitat suitability, where less suitable habitat would be costlier to move through (Eq. 1).

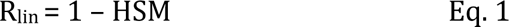

R_lin_ is the landscape resistance based on a linear transformation of habitat suitability (HSM) (Figure 3A). A nonlinear response would be, for example, when elephant movement across the landscape is only substantially impacted by highly unsuitable habitats (e.g., very high cost areas), but where medium- and low-cost areas do not impact movement. We used a negative exponential function to nonlinearly transform HSM to cost, R (Eq. 2).

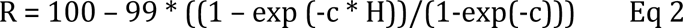

where R is resistance, H is suitability, and the factor c determines the shape of the curves. We generated resistance surfaces for c = 2 (slight nonlinear transformation) and c = 8 (pronounced nonlinear transformation) to allow for variable strengths of nonlinear responses (Figure 3B).

For each of the transformed resistance surfaces, we calculated three commonly used measures of landscape connectivity (McRae and Beier, 2007) in a pairwise fashion for 142 elephant sample locations for which we had microsatellite data for 9 nuclear DNA loci (SI Table 2). These connectivity measures included standard geographic Euclidean distance, Least-Cost Path (LCP; Adriaensen et al., 2003; Cushman et al., 2006), and resistance distance based on circuit theory (CT; McRae and Beier, 2007).

To validate connectivity models, cost distance is compared to genetic distance. We assumed an inverse relationship between gene flow and genetic distance (GD), where areas of high gene flow would result in low genetic distances among individuals and *vice versa*. Different estimates of GD vary in their ability to capture variation at the landscape scale (Shirk et al., 2012, 2017). We therefore quantified pairwise GD using four alternative GD estimates previously used in landscape genetic studies (Shirk et al., 2012; Kamvar et al., 2014; Milanesi et al., 2017; Tang et al., 2019) using the program R (R Core Team, 2019). Some GD estimates rely on the same data variables (e.g., allele presence and frequency) to quantify GD. The GD measures presented in this study are therefore not independent and there may be overlap between GD quantifications and model outcomes when GD estimates with highly similar data variables are used as response variables for model fitting. To represent response variables in our MLPE models, we calculated GD as 1) a value of 1 minus the proportion of shared alleles (D_PS_) using the “propShared” function in the R package “adegenet” (Jombart, 2008); 2) the number of allelic differences between two individuals using the “diss.dist” function in the R package “poppr” (Kamvar et al., 2014); 3) the Euclidean distance among a vector of allele frequencies using the “dist” function in the R package “adegenet”; 4) Reynolds’s distance with the “Reynolds.dist” function in the R package “poppr”.

Euclidean, LCP and CT distances represent the fixed effects in each model while the four quantifications of pairwise genetic distances among individuals represent the response variables. Using the “base” package in R (R Core Team, 2019), we scaled our response variables prior to fitting MLPE models to allow for the comparison with predictor variable values. The scale function in “base” centers and scales the columns of a numeric matrix (Becker et al., 1988). For each of three cost surface and four genetic distance response variables, we fitted MLPE models that considered as fixed effects Euclidean distance only, LCP and Euclidean distance, and CT and Euclidean distance, resulting in a comparison of 36 MLPE models in total. We include Euclidean distance as a fixed effect to our LCP and CT MLPE models to parse out geographic distance effects that are inherently included in LCP and CT (Row et al., 2017).

Model selection and fit was identified using Akaike’s information criterion, marginal R^2^ and conditional R^2^ values calculated for maximum likelihood population effects (MLPE) models where genetic distance is the response variable, resistance distances of connectivity models are fixed effects, and individual identification is used as random effects to account for pseudo-replication (Clarke et al., 2002; Van Strien et al., 2012). We fitted MLPE models between pairwise matrices of genetic distances and matrices of Euclidean, LCP or CT distances.

Akaike’s information criterion (AIC) was calculated in the “lme4” package in R (Burnham and Anderson, 2004; Bates et al., 2015), and marginal and conditional R^2^ were calculated in the R package “MuMIn” (Barton, 2009). The lowest AIC values indicate the MLPE model with the best balance between model fit and complexity (Burnham and Anderson, 2004), higher marginal R^2^ values represent higher predictive power of the fixed effects, whereas higher conditional R^2^ value represent higher total variance explained by the combined fixed and random effects (Edwards et al., 2008). Compared to traditional R^2^ values, marginal and conditional R^2^ values are favorable model selection criteria (Van Strien et al., 2012) because they do not necessarily increase with the addition of model parameters (Orelien and Edwards, 2008). Restricted maximum likelihood (REML) estimation of MLPE models has been recommended for determining accurate estimates of R^2^ (Verbeke and Molenberghs, 2000; Gurka, 2006), so we calculated conditional and marginal R^2^ values using REML. However, REML should not be used when comparing information criteria for MLPE models with different fixed effects, and we therefore did not include REML when we calculated AIC.

### Landscape connections

Out of the 36 MLPE models, we identified a single model that had the lowest AIC and the highest conditional and marginal R^2^. We independently compared AIC values per individual response variable since AIC values should not be compared across models that use different response variables. For each of our response variables, we identified the model with the lowest AIC.

Our final landscape connections map thus considers suitable habitats based on spatial data, validated by estimating gene flow as the inverse of genetic distance, to delineate areas in the landscape that may be important for maintaining population connectivity as part of elephant conservation initiatives. To create a summary map that considers both spatial and genetic data, we used Omniscape (McRae et al. 2016, Landau et al. 2021) to visualize habitat suitability using the top MaxEnt model (i.e., lowest AIC). Omniscape uses a moving window to estimate connectivity of surrounding cells in relation to one central cell using a CT approach, reducing the spatial bias introduced by genetic sampling locations (McRae et al., 2016; Landau et al., 2021) in the program Julia (Bezanson et al., 2017). The radius was set to 334 pixels (∼300km) to allow for connectivity mapping using a geographic scale relevant to both elephant space use (Loarie et al., 2009b; Roever et al., 2013; Huang et al., 2022) and gene flow (de Flamingh et al., 2018). We evaluated the Omniscape map by comparing Normalized Cumulative Current values at 10,000 locations where elephants were present (occurrence points) to 10,000 random locations (SI Figure 4A). The Normalized Cumulative Current represents only the impacts of the HSM on gene flow by accounting for the effects of geographic distances from the raw cumulative current map. We limited our evaluation extent so that it overlapped broadly with our genetic sampling extent, but excluded obvious unsuitable areas (e.g., the Indian Ocean; see SI Figure 4 for the evaluation extent boundaries). A two-sample z-test (Kitchens, 2002) was used to compare location points and random points using a custom script in R after confirming that the data were normally distributed.

**Figure 4.**
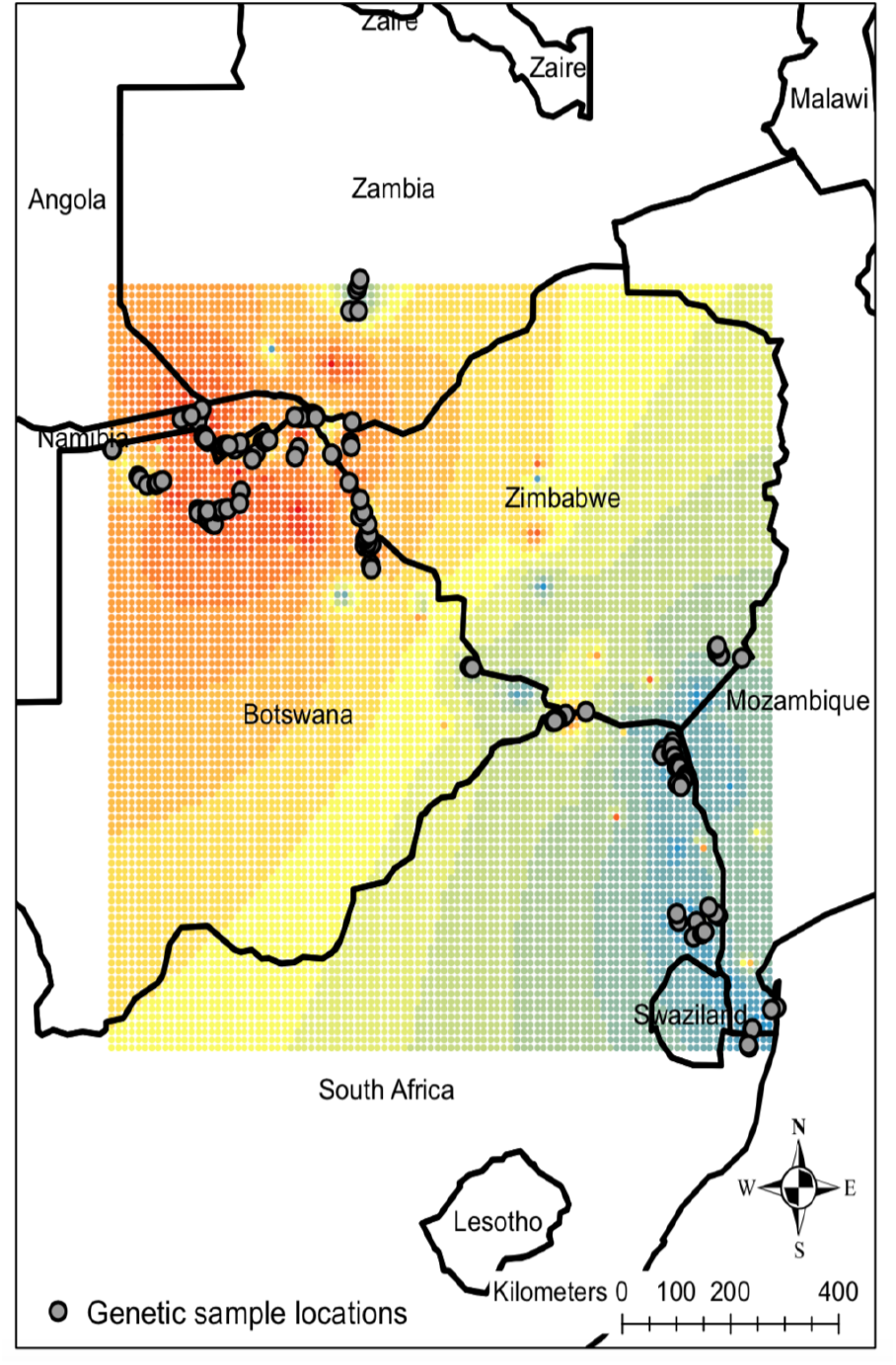
Visual representation of genetic patterns across the study extent, showing that areas in South Africa (e.g., Kruger National Park) were genetically similar (low interpolated genetic distances) relative to elsewhere. The interpolated genetic distance surface is a relative representation of gene flow across the study area and is dependent on the genetic variation captured by the marker system. Areas in blue indicate higher relative gene flow (lower genetic distance) and areas in red indicate lower relative gene flow (higher genetic distance) among 142 elephant genetic samples (grey circles).

## RESULTS

### Elephant occurrence data

From the Conservation Ecology Research Unit, occurrence data from 80 elephants were used, of which 65 were from elephant breeding herds and 15 were male elephants. Our spatial database included a total of 53,954 location points. The temporal scale of data varied per individual since GPS collars were deployed in different years, but overall, the elephant monitoring occurred from 2002 to 2015 across all seasons.

### Elephant genetic data

We genotyped 142 elephant fecal samples for 9 nuclear DNA microsatellite loci developed in elephants (SI Table 1;Comstock et al., 2000; Eggert et al., 2000; Archie et al., 2003). Across the elephants used in this study, the loci were highly variable, with the number of alleles ranging between six and thirteen per locus (mean = 8.89). The observed heterozygosity was slightly lower than the expected heterozygosity (SI Table 1), likely reflecting a limited degree of population differentiation among populations across Southern Africa.

### Habitat suitability

We found no significant correlation (Pearson’s R <0.5, p >0.5) between our MaxEnt environmental layers when including “ambient population” distribution as a measure of human pressure (SI Figure 5). In contrast, the transformed gradient-based human distance metric was correlated with our GSW environmental variable (r > 0.60; Figure 2). Also, the gradient-based distance from high human densities performed poorly compared to ambient population (Table 1). We therefore used raw ambient human population data (Figure 2E) rather than transformed gradient-based distance data to represent human population distribution in our MaxEnt models.

**Figure 5.**
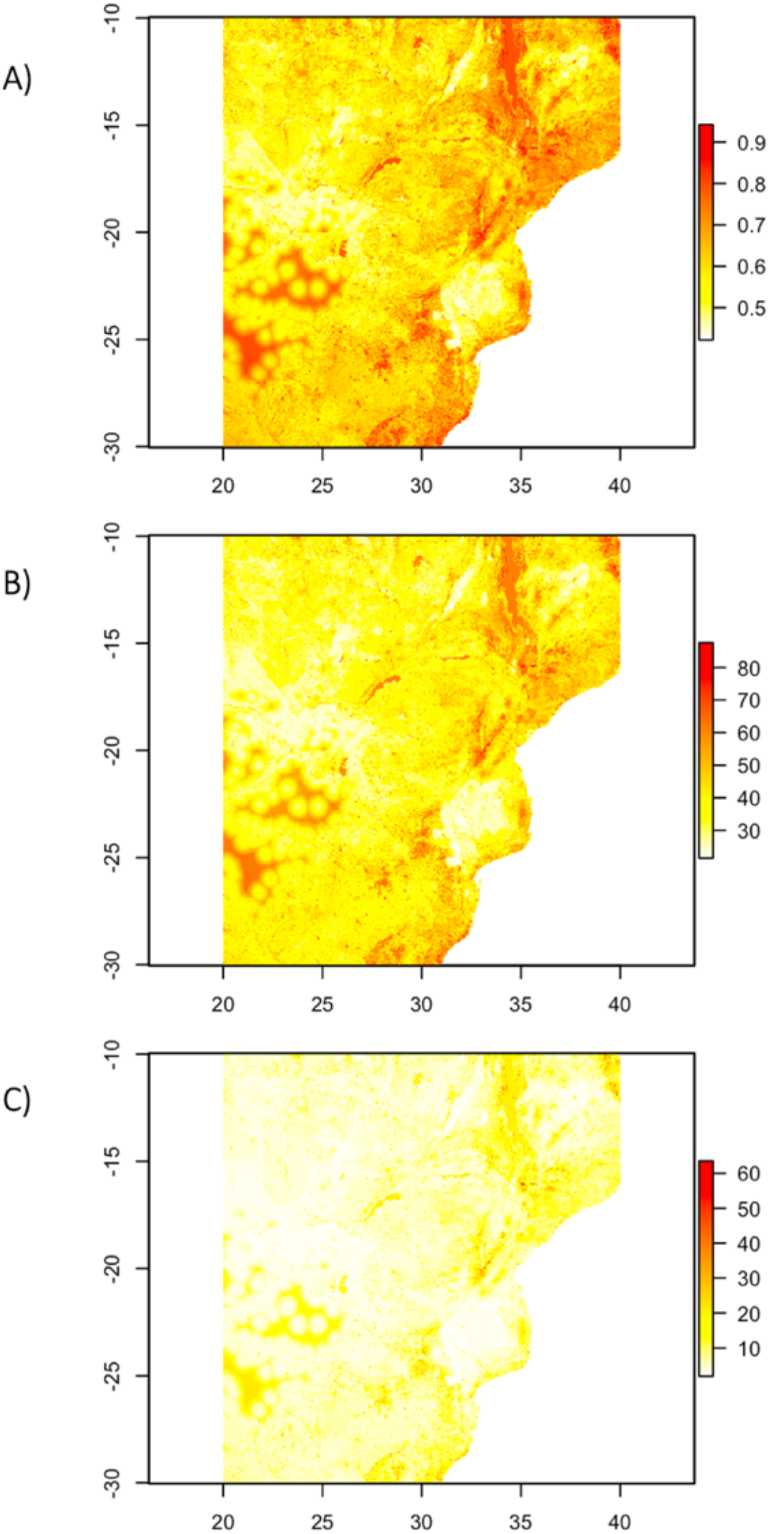
Transformation of the MaxEnt habitat suitability model into three alternative representations of landscape resistance. The linear transformation (panel A) assumes that resistance is inversely related to habitat suitability, while the nonlinear transformations allowed for slight nonlinear responses (panel B) and pronounced nonlinear responses (panel C) to unsuitable habitats. The X and Y axes indicate longitude and latitude, and map colors indicate areas in the landscape that are very costly (red) to less costly (white) for elephants to move through.

We tested four regularization beta parameters in Maxent (*β* = 1, 2, 3, 4) for each of the background datasets (range-based and use-based background). The range-based background dataset resulted in slightly higher AUC than use-based background dataset and we therefore selected the range-based dataset. We found that *β* = 1 had the highest regularization gain and AUC, and we therefore selected the MaxEnt model calculated using range-based background data and *β* = 1 as the top habitat suitability model (SI Figure 6).

We investigated how each of the environmental variables contributed to the predicted habitat suitability model and found that human density as “ambient population” distribution, distance to water, and slope contributed in a non-linear fashion. All three of these environmental variables decreased in their habitat suitability until they reach a threshold beyond which they were not suitable for elephants (SI Figure 7). The habitat suitability model indicated elephants occur in areas of low human density, close to water, and with no or low slopes. Intermediate values of primary productivity contributed the most to our predictive habitat suitability model, which is congruent with the distribution of elephants in our study area that mostly occurs in landscapes with intermediate productivity (e.g., savanna or non-woody habitats; Mapaure and Campbell, 2002; Young et al., 2009a).

### Spatial visualization of gene flow

Using GD calculated in AIS, we found no correlation between genetic and geographic distances (partial Mantel test r = -0.035, probability of observing a ≥ correlation P = 0.9, probability of observing a ≤ correlation P = 0.06). To visualize genetic patterns across the landscape, we interpolated genetic distances to form a landscape shape in the program AIS. The interpolated GD surface is a relative representation of gene flow across the study area because the GD estimates are relative to genetic variation captured by the microsatellite marker system. Our GD surface indicated there was less constrained gene flow in the south-eastern region of the study area than in the north-western region (Figure 4).

### Landscape genetics

We linearly and nonlinearly transformed the top MaxEnt HSM (range-based background points, *β* = 1) into three alternative representations of landscape resistance (Figure 5). The MLPE model assessment consistently identified CT distances calculated for the pronounced nonlinear transformation (c = 8) as the connectivity measure that best explained the observed gene flow for all four genetic metrics across the landscape (Table SI 2). These genetic metrics are not independent and will likely result in similar delineation and visualization of landscape connections. However, to limit replicate analyses we focused on one genetic metric, D_PS_, which has often been used in landscape genetic studies (e.g., see Hazlitt et al. (2004) and Milanesi et al. (2017) for examples of usage of D_PS_ for model evaluation and Waits and Storfer (2015) for a general overview of D_PS_ as a genetic metric).

### Landscape connections

Because we identified CT distances as the connectivity measure that best explained the gene flow across our landscape, we used CT distances in Omniscape (Shah and McRae, 2008) to delineate and visualize areas that represent landscape connections for elephants (Figure 6A). We found areas in our landscape connections map where the impact of different environmental determinants was noticeable; primary productivity (Figure 6B) and “ambient human population” distribution (Figure 6C) seem to be the primary drivers of connectivity in parts of the landscape. The mean Normalized Cumulative Current at elephant occurrence points was higher than the mean at random points (*z* = -78.952, *p* < 0.001), indicating the connectivity map generated using Omniscape was a good predictor of where elephants occur (SI Figure 4B).

## DISCUSSION

Here we provide a new framework for empirically delineating landscape connections for species by integrating habitat suitability and gene flow. We found a pronounced nonlinear response to habitat suitability that suggests elephant movement and gene flow are mostly impacted by very unsuitable habitats, and that moderately unsuitable habitat impedes connectivity between elephant populations to a lesser degree. This nonlinear response agrees with space use described for some elephant populations in our study region; for example, elephants in Botswana are known to range across populated areas to access food and water (Hoare and Du Toit, 1999; Jackson et al., 2008).,Despite this, we identified ambient human population as a likely a barrier to elephant habitat suitability and there is likely a population threshold that eventually prevents gene flow. However, the challenges faced by elephants in moving across a landscape may vary for different populations (Huang et al., 2022), requiring consideration of different factors when analyzing paths of connectivity for different sets of populations.

The pronounced nonlinear response to habitat suitability agrees with criticisms of typical habitat suitability models used in gene flow analyses (Keeley et al. 2017) as the linear model performed poorly. However, the performance of the pronounced non-linear transformation of habitat suitability to resistance indicates that habitat suitability models can still provide insight to gene flow across complex landscapes. Specifically, the resistance map using transformed suitability identified areas with different limiting factors (ambient human population and vegetation) to gene flow. Through this method, empirical models can be created and used to determine gene flow (but see section “Caveats and future directions”).

The analyses also indicated that landscape representations that consider multiple paths of connectivity (isolation by resistance based on circuit theory; IBR) perform better at predicting gene flow across the landscape than singular paths (LCP) or geographic distance alone. Similar IBR frameworks have been more effective at explaining gene flow for black bears (Ursus americanus, Cushman et al., 2006), hedgehogs (Erinaceus europaeus, Braaker et al., 2017), and other plant and animal species (McRae and Beier, 2007).

Our landscape connections map in Omniscape (Figure 6) shows that gene flow and connectivity appear to be influenced by different environmental variables across the landscape, where connectivity in some areas is clearly driven by a single variable. For example, primary productivity as a proxy of food availability was the primary driver of connectivity in regions surrounding the Makgadikgadi pans in Botswana (Figure 6B). The low primary productivity associated with the pans is reflected as low predicted connectivity or suitability of this area as a landscape connection. We found human density as the primary driver of connectivity in the landscape that links elephants from Botswana with elephants in South Africa in the areas adjacent to northern parts of Kruger National Park (Figure 6C), which is also consistent with the results of Huang et al., (2022) using a more extensive spatial dataset. Although not highlighted in Figure 6, examples of the impacts of other environmental variables include slope as the primary driver of landscape connectivity for regions that overlap with the Lebombo mountain range (du Toit, 1929; Saggerson and Bristow, 1983) that separates elephants in southern Kruger National Park (South Africa) and Maputo Special Reserve (Mozambique). Similarly, distance to water was the primary driver of connectivity in the Caprivi region of Namibia.

When demarcating landscape connections for conservation, researchers and conservation stakeholders should therefore consider that landscape connections may be dependent on the specific geographic area and environmental variables under consideration, and that a “one-map-fits-all” approach should be avoided. Huang et al. (2022) provide case studies of different elephant populations that each have unique conservation challenges, and also describe ways to assess whether landscape connectivity could be restored if challenges are negated, or whether such efforts would be futile. We therefore urge researchers and conservation stakeholders to consider additional factors other than landscape connectivity and gene flow when demarcating landscape connections for conservation. Factors such as land ownership (Pinter-Wollman, 2012), human-elephant conflict mitigation (Jackson et al., 2008; Pinter-Wollman, 2012), the direct and indirect impact of creating landscape connections on local indigenous communities (Baldus et al., 2007), landscape connections overlapping with poaching hotspots (Zafra-Calvo et al., 2018; Schlossberg et al., 2019), and other sociopolitical factors need to be integrated into conservation decisions.

### Caveats and future directions

Our goal was to test and validate a general methodology that could be replicated across systems. However, elephant space use and habitat suitability modeling may be influenced by factors that were not considered in this study. For example, the pronounced nonlinear response to occurrence-based habitat suitability models suggests that alternative or expert-based quantifications of habitat use might be needed to accurately establish elephant connectivity requirements. Researchers could, for example, separate elephant movements into different behavioral states using hidden Markov models fitted in a Bayesian framework (Leos-Barajas and Michelot, 2018; Wang, 2019; Vogel et al., 2020), and identify from those states the environmental variables that are important for establishing or maintaining connectivity. For example, Keeley et al., (2017) showed that habitat suitability is a poor proxy for landscape connectivity during dispersal and mating movements in kinkajous (*Potos flavus*), where tolerance for unsuitable habitat during dispersal seems common. Mateo-Sánchez et al., (2015) showed that dispersing Cantabrian brown bears (*Ursus arctos*) might be more flexible in their dispersal movement behavior than they are in their habitat use behavior. Determining behavioral states may be especially relevant to elephant landscape connections because exploratory movements within corridors may be fast and directional compared to encamped foraging movements that are slow and meandering (Vogel et al., 2020). It may therefore be beneficial for future studies to consider both behavior and space use when delineating landscape connections for African elephants.

In addition to behavior, other aspects not considered here could also impact the accurate delineation of landscape connections. Mashintonio et al., (2014) showed elephants select habitat based on environmental qualities at multiple spatial scales, and it may therefore be informative to incorporate habitat suitability modeling at different scales. Researchers could, for example, use a Gaussian pixel smoothing algorithm approach, which can be effective for determining the scale at which elephants select resources (Mashintonio et al., 2014).

In addition to scale and resolution, seasonality and sex are crucial drivers of elephant dispersal and land use patterns (Young et al., 2009b). Future studies could consider seasonal and sex-specific differences when predicting gene flow across the landscape by generating individually modeled landscape connections maps for males and females, and for wet and dry seasons (Young et al., 2009a; Purdon et al., 2018). Sex- and season-based landscape connections maps may be especially important for elephant connectivity mapping since dispersal is predominantly male-mediated (Nyakaana and Arctander, 1999; Roca et al., 2005), and males typically have larger home ranges and disperse farther than females with young calves (Mole et al., 2016). For conservation, protecting dispersal and movement of males may be a critical component for maintaining adequate gene flow across the region.

Despite these limitations, the conservation genetic approach developed and applied here integrates multidisciplinary datasets and methods in an innovative way and may therefore be a useful framework for future studies on elephant and other taxa that aim to develop spatially and genetically informed conservation strategies. In particular, an habitat suitability models robust to integrating data types. Although we applied these methods to a robust dataset, MaxEnt and AIS are effective even with small sample sizes, and this approach should be applicable to a range of taxa, especially taxa with depauperate occurrence data.

### Broader context and application

The defragmentation of conservation areas through the development and maintenance of landscape connections could induce regional demographic stability in elephant numbers, increase available areas to roam thus allowing for space use that may change seasonally, reduce local impact that elephants have on the landscape (van Aarde and Jackson, 2007; Huang et al., 2022), and mitigate the consequences of genetic isolation by promoting gene glow and increasing genetic diversity (Allendorf and Luikart, 2009; Burkart et al., 2016; Orton et al., 2020). Integrating multifaceted landscape habitat modeling with genetic analyses has been proposed and applied as a conservation tool, and it is an expedient method through which landscape connections hypotheses can be tested. For example, the integration of spatial and genetic analyses was used to detect and evaluate landscape connectivity and movement corridors for wolves (Canis lupus, Kabir et al., 2017), Cantabrian brown bears (Mateo-Sánchez et al., 2015), rodents (Wang et al., 2008) and birds (Klinga et al., 2019). In this study, we show that the integration of spatial landscape modeling and genetic analyses can also be used to delineate landscape connections for African elephant conservation planning.

## ACKNOWLEDGEMENTS

We dedicate this paper to the memory of Rudi van Aarde whose life-long passion and dedication to conservation was and continues to be a driving force for the protection of Africa’s elephants. We thank Camilla Nørgaard for permission to include the late Rudi van Aarde as co-author. For funding we thank the International Fund for Animal Welfare; the Conservation Ecology Research Unit (CERU) of the University of Pretoria; the Conservation Foundation (Zambia); the US Fish and Wildlife Service African Elephant Conservation Fund. In addition to support from these sources, AdF received a Francis M. and Harlie M. Clark Research Support Grant, a Harley J. Van Cleave Research Award, a University of Illinois Graduate College Dissertation Project Travel Grant, and support from the Cooperative State Research, Education, and Extension Service, US Department of Agriculture under project number ILLU 875–952. For facilitating sample collection, we thank Elephant Without Borders (Botswana) and the CERU team. The Zambian Wildlife Authority, the Department of Wildlife and National Parks (Botswana), South African National Parks (SANParks) and the Department of Agriculture, Forestry and Fisheries (South Africa) sanctioned the research. We acknowledge technical support provided by the University of Pretoria’s Sequencing Facility; the High-Throughput Sequencing and Genotyping Unit of the University of Illinois at Urbana-Champaign (UIUC); and the Illinois Campus Cluster, a computing resource that is operated by the Illinois Campus Cluster Program (ICCP) in conjunction with the National Center for Supercomputing Applications (NCSA) and which is supported by funds from UIUC.

## ETHICS STATEMENT

The Animal Ethics Committee of the University of Pretoria (AUCC-040611-013) and the Botswana Ministry of Environment, Wildlife, & Tourism (OP 46/1 LXXXV 89) reviewed and approved the telemetry collaring of the elephants. Collection and export/import of elephant dung was sanctioned by appropriate authorities prior to collection, including South African National Parks, the Department of Agriculture, Forestry and Fisheries in South Africa and the United States Department of Agriculture in the United States of America.

## AUTHOR CONTRIBUTION

AdF, RLS, RJvA and ALR conceived and developed the methodological analysis approach. AdF, TINPS and CD completed the molecular work, and AdF and NA conducted the computational analyses. AdF wrote the original draft of the manuscript, all authors provided critical feedback throughout the process of interpretation of the data and of manuscript preparation, and approved of the final version of the manuscript.

## DATA AVAILABILITY

All bioinformatic code associated with this manuscript may be found at https://github.com/adeflamingh/de_Flamingh_et_al_Landscape_Connectivity Genetic data is available on DRYAD: XXXXX

