## Supplementary Files for "Integrating habitat suitability modeling with gene flow improves delineation of landscape connections among African savanna elephants"

### Supplementary information:

#### Supplementary Tables

SI Table 1. Marker names, profiles and characteristics^4^ of nine microsatellite loci used in this study.

| **Marker**  **Name** | **Locus Name** | **Repeat Motif** | **Size (bp)** | **Ta-°C** | **No of alleles** | **HO** | **HE** |
| --- | --- | --- | --- | --- | --- | --- | --- |
| **LaT08^2^** | 1 | (TAGA)16 | 166–234 | 56 | 13 | 0.81955 | 0.86347 |
| **Lat13^1^** | 2 | (CATC)21 | 173–262 | 56 | 10 | 0.62963 | 0.76649 |
| **Lat17^1^** | 3 | (GGAT)15… (GGAT) | 324–352 | 56 | 8 | 0.71212 | 0.80626 |
| **Lat24^1^** | 4 | (GGAT)22 | 203–231 | 56 | 8 | 0.77953 | 0.84075 |
| **FH1^2^** | 5 | (CA)12 | 81-89 | 55 | 6 | 0.56522 | 0.61660 |
| **FH39^2^** | 6 | (CA)18 | 232-256 | 60 | 12 | 0.63704 | 0.78441 |
| **FH102^2^** | 7 | (CT)11(CA)14 | 175-187 | 60 | 6 | 0.51773 | 0.49862 |
| **LA5^3^** | 8 | (CA)13 | 139–153 | 52 | 8 | 0.56028 | 0.55996 |
| **Lat25^1^** | 9 | (CCAT)15 | 287–321 | 52 | 9 | 0.66923 | 0.83710 |
| **Mean** |  |  |  |  | 8.889 | 0.65448 | 0.73041 |
| **s.d.** |  |  |  |  | 2.421 | 0.10179 | 0.13554 |

^1^ Comstock et al., 2000

^2^ Archie et al., 2003

^3^ Eggert et al., 2000

^4^ Number of alleles per locus, observed heterozygosity (HO) and expected heterozygosity (HE)

SI Table 2. Calculated MLPE models for three resistance surfaces (linear, slight nonlinear, and pronounced nonlinear transformation of HSM) and four genetic distance response variables (D_PS_ = 1 minus the proportion of shared alleles; GD_Aldiff = genetic distance as the number of allelic differences between two individuals; GD_euc = genetic distance as the Euclidean distance among a vector of allele frequencies; GD_TotGD = Reynold’s genetic distance measure). For each of these resistance surface and genetic distance combinations, we fitted MLPE models that considered as fixed effects geographic Euclidean distance (Geo ED) only, Least-cost path distance (LCP) and Geo ED, and resistance distance (Circuit Theory) and Geo ED, resulting in a comparison of 36 MLPE models in total. Based on AIC, conditional and marginal R^2^, that the model with CT and Geo ED calculated using the pronounced nonlinear HSM transformation is the best predictor across genetic markers. D_PS_ was used for visualization of gene flow and landscape connections across the landscape (indicated in **boldface**).

|  | Linear Transformation | | | Slight nonlinear transformation | | | Pronounced nonlinear transformation | | |
| --- | --- | --- | --- | --- | --- | --- | --- | --- | --- |
|  | AIC | R2m | R2c | AIC | R2m | R2c | AIC | R2m | R2c |
| D_PS_ ~Geo ED | -30950.4 | 0.001 | 0.099 | -30950.4 | 0.001 | 0.099 | -30950.4 | 0.001 | 0.099 |
| D_PS_ ~LCP + Geo ED | -30954.4 | 0.002 | 0.102 | -30953.4 | 0.002 | 0.101 | -30954.1 | 0.002 | 0.102 |
| D_PS_ ~CT + Geo ED | -30968.3 | 0.004 | 0.103 | -30987.4 | 0.007 | 0.107 | **-31006.6** | **0.010** | **0.111** |
| GD_Aldiff ~Geo ED | 26920.6 | 0.001 | 0.099 | 26920.6 | 0.001 | 0.099 | 26920.6 | 0.001 | 0.099 |
| GD_Aldiff ~LCP + Geo ED | 26916.6 | 0.002 | 0.102 | 26917.6 | 0.002 | 0.101 | 26916.9 | 0.002 | 0.102 |
| GD_Aldiff ~CT + Geo ED | 26902.7 | 0.004 | 0.103 | 26883.7 | 0.007 | 0.107 | 26864.4 | 0.010 | 0.111 |
| GD_euc ~Geo ED | 113.5 | 0.001 | 0.095 | 113.5 | 0.001 | 0.095 | 113.5 | 0.001 | 0.095 |
| GD_euc ~LCP + Geo ED | 109.5 | 0.002 | 0.097 | 110.6 | 0.002 | 0.097 | 110.0 | 0.002 | 0.097 |
| GD_euc ~CT + Geo ED | 99.9 | 0.003 | 0.098 | 81.3 | 0.007 | 0.107 | 60.4 | 0.009 | 0.106 |
| GD_TotGD ~Geo ED | -46927.4 | 0.001 | 0.092 | -46927.4 | 0.001 | 0.092 | -46927.4 | 0.001 | 0.092 |
| GD_TotGD ~LCP + Geo ED | -46931.3 | 0.002 | 0.094 | -46930.2 | 0.002 | 0.094 | -46930.8 | 0.002 | 0.094 |
| GD_TotGD ~CT + Geo ED | -46938.7 | 0.003 | 0.094 | -46956.8 | 0.006 | 0.098 | -46978.3 | 0.009 | 0.103 |

#### Supplementary Figures

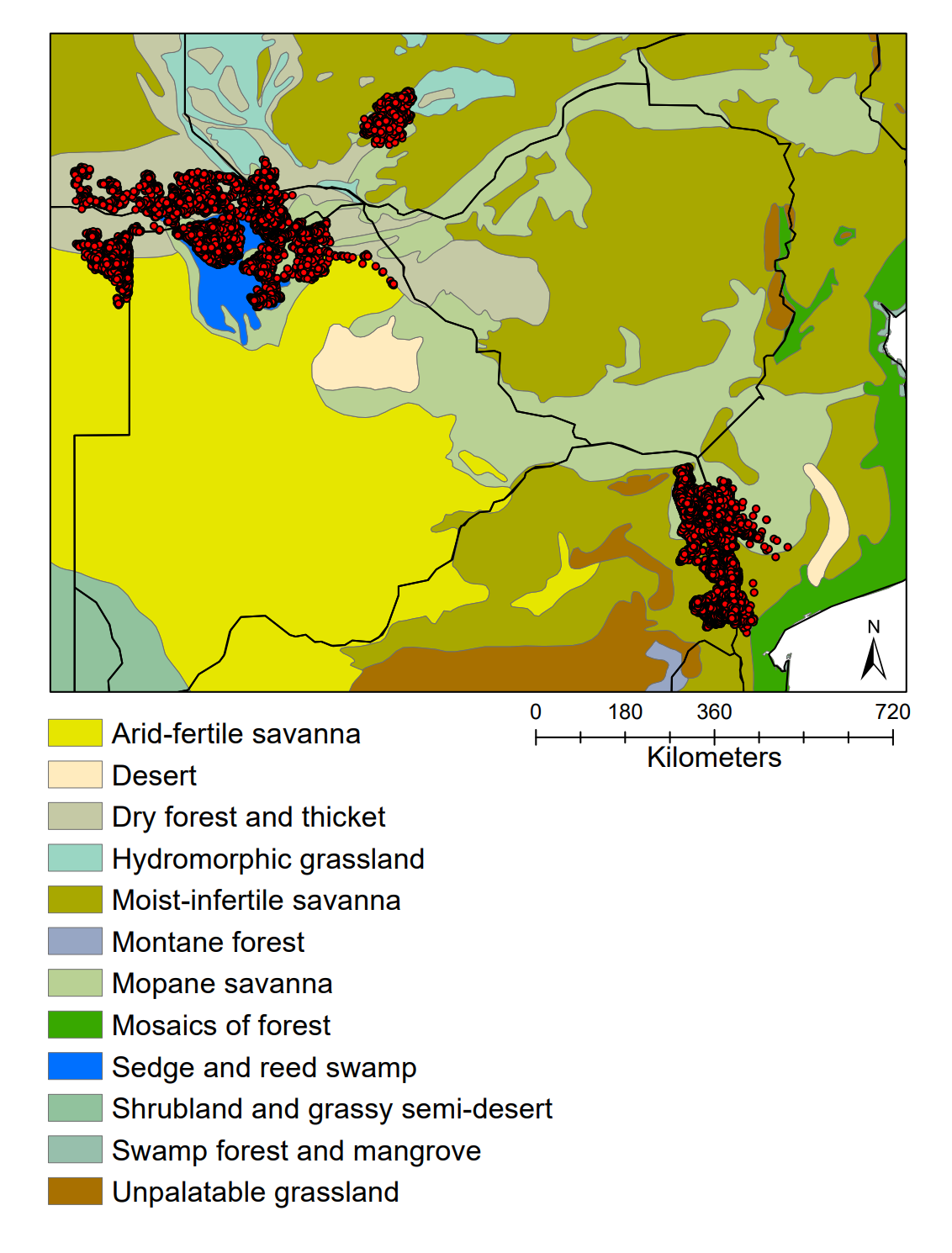

SI Figure 1. The study area includes seven Southern African countries (delineated by black lines) namely Angola, Botswana, Mozambique, Namibia, South Africa, Zambia and Zimbabwe. Elephant occurrence data (red dots) spans a range of different vegetation classes (White, 1983) across the study area.

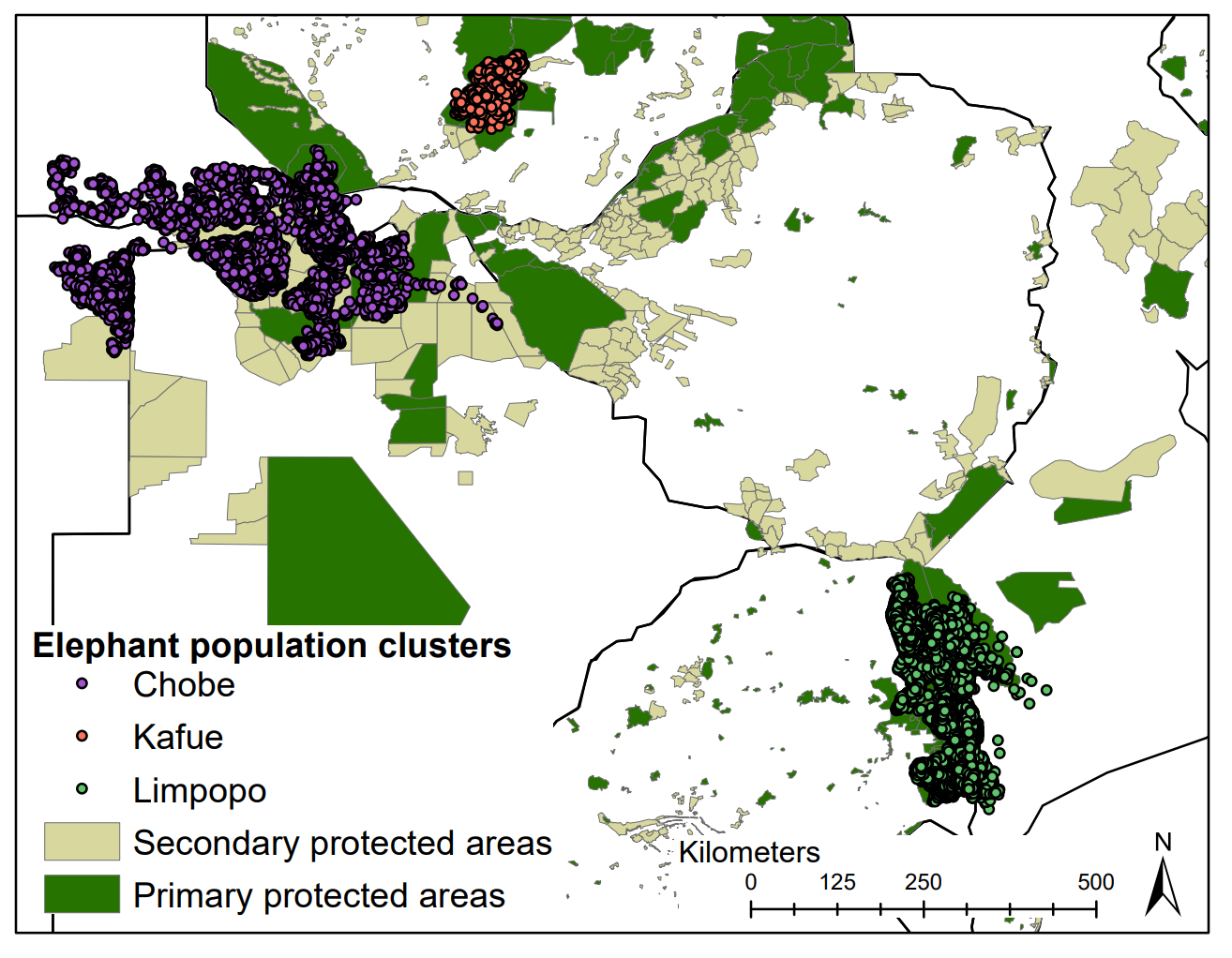

SI Figure 2. Spatial location data from 80 elephants, forming part of 3 regional population clusters that include the Chobe, Kafue, and Limpopo elephant population clusters.

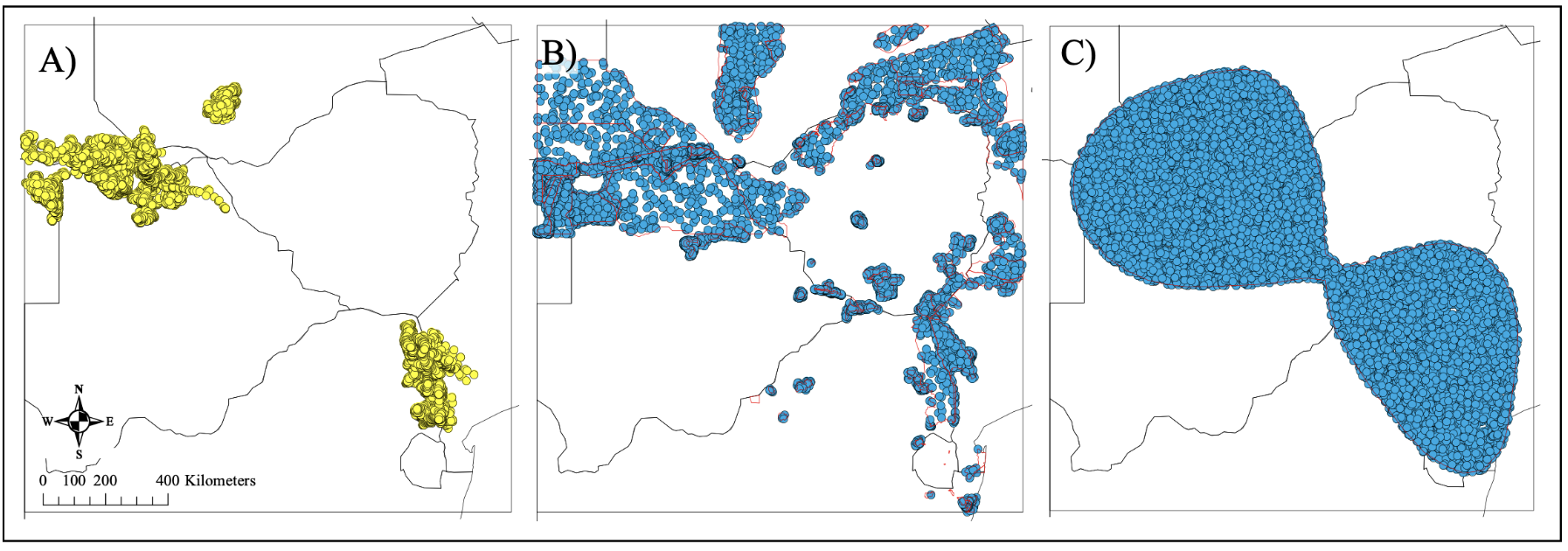

SI Figure 3. MaxEnt compares environmental conditions at background location data points to environmental conditions at known presence (occurrence) locations (panel A – yellow points). Background data should reflect the environmental conditions that are relevant to the species for which the model is generated. We tested two different background datasets: background data generated for areas of known elephant range (range-based) as demarcated by the IUCN Red List of Threatened Species (panel B – red lines indicate known range from Blanc, (2008)); and background data from areas included in an 80% convex hull of presence points (panel C – red lines indicate 80% convex hull). For each of these datasets we generated 10 000 random points (shown in blue in panels B and C) to compare to occurrence data.

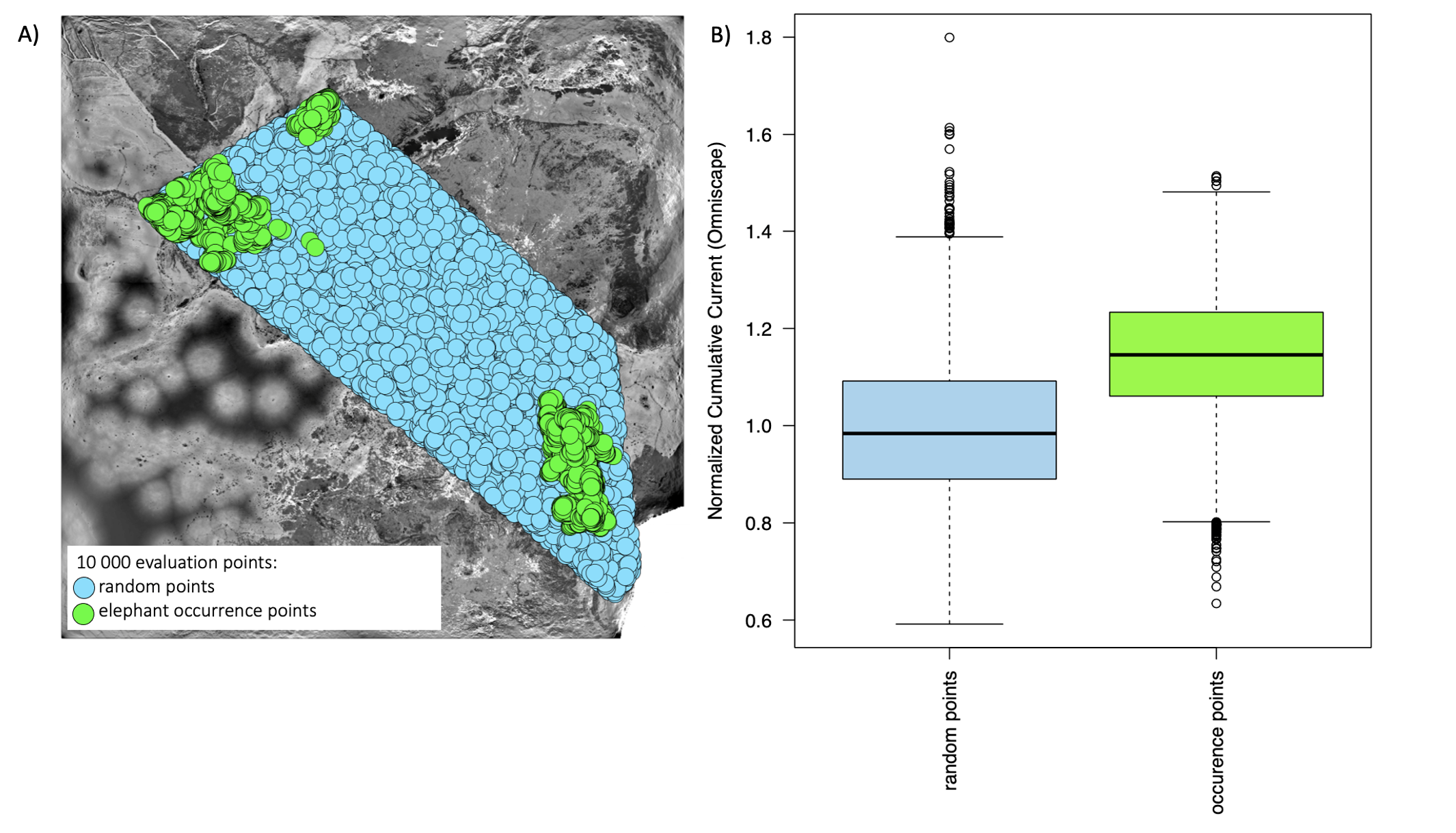

SI Figure 4. Omniscape LC map evaluation, which compared Normalized Cumulative Current values at random points (panel A - blue) to values at elephant occurrence points (panel A - green). A two-sample z-test (Kitchens, 2002) showed that the mean Normalized Cumulative Current at elephant occurrence points was significantly higher than the mean at random points (z = -78.952, p-value < 0.0001, panel B), indicating that the Omniscape landscape connections map has more current, or net random walks at elephant occurrences.

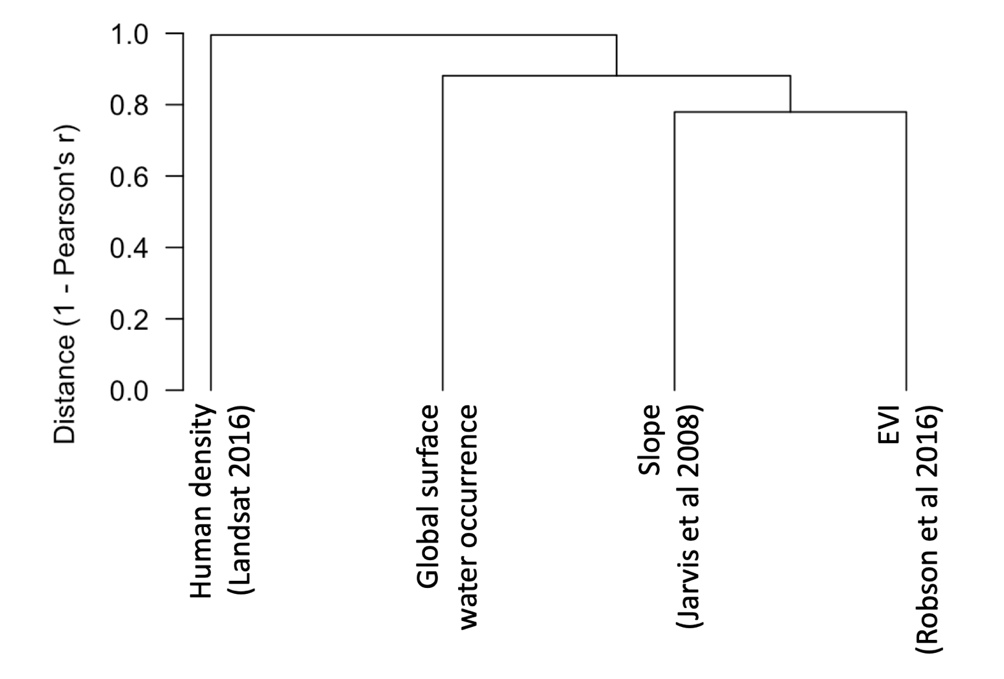

SI Figure 5. There was no significant correlation between any of the environmental variables included in the MaxEnt model when using “ambient population” distribution as our human density indicator. We tested Pearson’s R cut-off values of 0.7 (which is the least stringent and allows for most collinearity), 0.6, and 0.5 (the most stringent, presented here) and found no significant collinearity for any of the values.

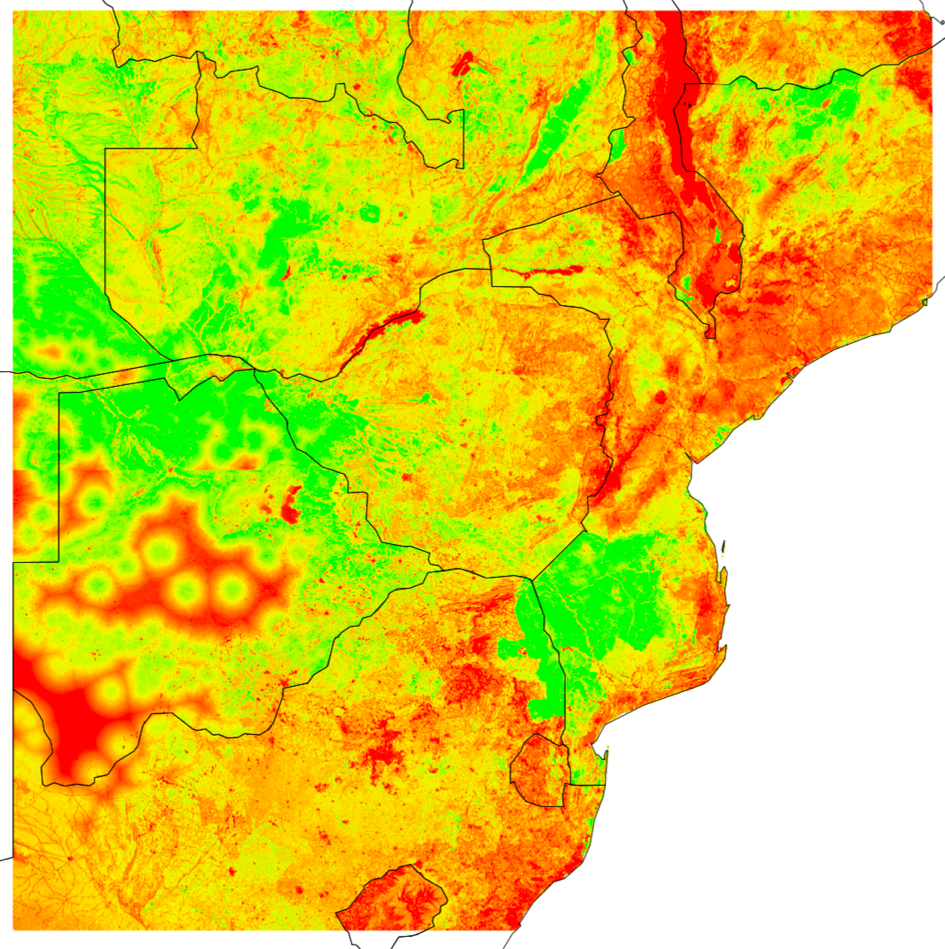

SI Figure 6. The predicted habitat suitability model (HSM) using MaxEnt with range-based background data and a regularization betaparameter of 1. Suitable elephant habitat is shown in green while less suitable habitats are shown in shades of yellow and red.

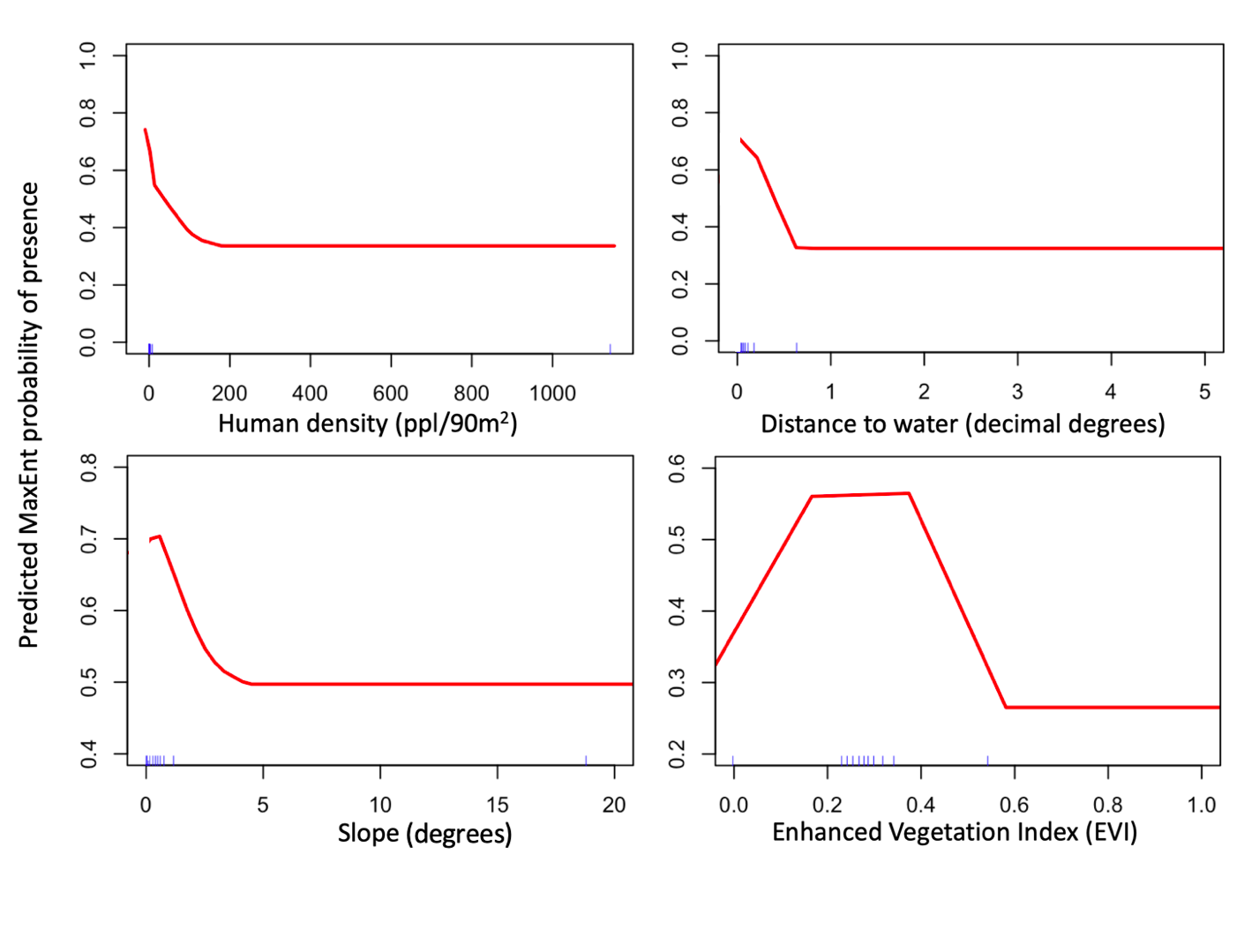

SI Figure 7. Response curves from the top MaxEnt habitat suitability model with human density as “ambient population” presence. Distance to water and slope are suitable for elephants at low environmental values (low human densities, close to water, and habitats with low slopes), and all three of these environmental variables decreased in their habitat suitability until they reached threshold environmental values beyond which they were not suitable habitats for elephants. Intermediate values of primary productivity (Enhanced Vegetation Index) contributed the most to predicted habitat suitability.
